# Balancing selection at a wing pattern locus is associated with major shifts in genome-wide patterns of diversity and gene flow

**DOI:** 10.1101/2021.09.29.462348

**Authors:** María Ángeles Rodríguez de Cara, Paul Jay, Quentin Rougemont, Mathieu Chouteau, Annabel Whibley, Barbara Huber, Florence Piron-Prunier, Renato Rogner Ramos, André V. L. Freitas, Camilo Salazar, Karina Lucas Silva-Brandão, Tatiana Teixeira Torres, Mathieu Joron

**Author notes:** contributed equally (María Ángeles Rodríguez de Cara, Paul Jay, Quentin Rougemont).

## Abstract

Selection shapes genetic diversity around target mutations, yet little is known about how selection on specific loci affects the genetic trajectories of populations, including their genome-wide patterns of diversity and demographic responses. Here we study the patterns of genetic variation and geographic structure in a neotropical butterfly, *Heliconius numata*, and its closely related allies in the so-called melpomene-silvaniform clade. *H. numata* is known to have evolved an inversion supergene which controls variation in wing patterns involved in mimicry associations with distinct groups of co-mimics whereas it is associated to disassortative mate preferences and heterozygote advantage at this locus. We contrasted patterns of genetic diversity and structure 1) among extant polymorphic and monomorphic populations of *H. numata*, 2) between *H. numata* and its close relatives, and 3) between ancestral lineages. We show that *H. numata* populations which carry the inversions as a balanced polymorphism show markedly distinct patterns of diversity compared to all other taxa. They show the highest genetic diversity and effective population size estimates in the entire clade, as well as a low level of geographic structure and isolation by distance across the entire Amazon basin. By contrast, monomorphic populations of *H. numata* as well as its sister species and their ancestral lineages all show lower effective population sizes and genetic diversity, and higher levels of geographical structure across the continent. One hypothesis is that the large effective population size of polymorphic populations could be caused by the shift to a regime of balancing selection due to the genetic load and disassortative preferences associated with inversions. Testing this hypothesis with forward simulations supported the observation of increased diversity in populations with the supergene. Our results are consistent with the hypothesis that the formation of the supergene triggered a change in gene flow, causing a general increase in genetic diversity and the homogenisation of genomes at the continental scale.

## Introduction

Genetic diversity is shaped by selective processes such as stabilizing or disruptive selection, and by demographic processes such as fluctuations in effective population size. Empirical studies on genetic diversity within and among populations abound, fuelled by an increasing availability of whole genome data, and spurred by our interest in understanding the underlying causes of variation in diversity (e.g. Beichmann 2018, Muers 2009; Murray 2017; Nielsen et al. 2009). At the locus scale, strong directional or disruptive selection tends to reduce diversity within populations (Mitchell-Olds et al. 2007), while balancing selection tends to enhance diversity (Charlesworth 2006). Genome-wide factors reducing diversity include low effective population sizes, generating drift, while high genetic diversity is enhanced by large population sizes and gene flow. Overall, it is well recognised that demographic changes should have a genome-wide effect on diversity, while positive selection is expected to play a role on the sites within and around the genes involved in trait variation (Glinka et al. 2003, Muers 2009, Nielsen et al. 2009).

Variation in behaviour and life-history traits, for instance involving changes in offspring viability or dispersal distance, may also affect species demography, and thus whole genome genetic diversity. However, whether and how genetic variability in a population may be driven by phenotypic evolution at certain traits is poorly understood, and confounding effects may affect patterns of genomic diversity, such as variation in census population size or colonization history. Dissecting how selection on a trait may affect genome-wide diversity can be tackled by comparing closely-related populations differing at this trait coupled with knowledge of when the differences evolved. Here, we took advantage of the dated introgressive origin of a chromosomal inversion associated with major life-history variation to study the demographics and whole genome consequences of changes in the selection regime at a major-effect locus.

*Heliconius* butterflies are aposematic, chemically-defended butterflies distributed over the American tropics from Southern Brazil to Southern USA (Emsley 1965; Brown 1979) (**Fig 1A**). *Heliconius* butterflies are well-known for visual resemblance among coexisting species, a relationship called Müllerian mimicry which confers increased protection from bird predators through the evolution of similar warning signals (Sheppard et al. 1985). Most species are locally monomorphic, but their mimicry associations vary among regions, and most species display a geographic mosaic of distinct mimetic “races” through their range. In contrast to most *Heliconius* species, the tiger-patterned *Heliconius numata* is well-known for maintaining both mimicry polymorphism within localities, with up to seven differentiated coexisting forms, and extensive geographic variation in the distribution of wing phenotypes (Brown & Benson 1974; Joron et al. 1999). Forms of *H. numata* combine multiple wing characters conveying resemblance to distinct sympatric species in the genus *Melinaea* and other local Ithomiini species (Nymphalidae: Danainae). Polymorphism in *H. numata* is controlled by a supergene, *i*.*e*. a group of multiple linked functional loci segregating together as a single Mendelian locus, coordinating the variation of distinct elements of phenotype (Brown & Benson 1974; Joron et al. 2006). Supergene alleles are characterized by rearrangements of the ancestral chromosomal structure, forming three distinct chromosomal forms with zero (ancestral type, Hn0), one (Hn1) or three chromosomal rearrangements (Hn123) (**Fig 1B**). The ancestral arrangement, Hn0, devoid of inversions, is fixed in most *Heliconius* species (although an inversion in the same region evolved independently in a distantly-related *Heliconius* lineage (Edelman et al. 2019)). Arrangement Hn1 contains a 400kb inversion called P_1_ originating from an introgression event about 2.2 My ago from *H. pardalinus*, in which P_1_ is fixed (Jay et al. 2018). This introgression is thought to be the founding event triggering the formation of the supergene and the maintenance of polymorphism in *H. numata* (Jay et al. 2018). Arrangement Hn123 displays two additional inversions, P_2_ and P_3_, in linkage with P_1_, and therefore originated after the introgression of P_1_ into the *H. numata* lineage (Jay et al. 2021).

**Figure 1.**
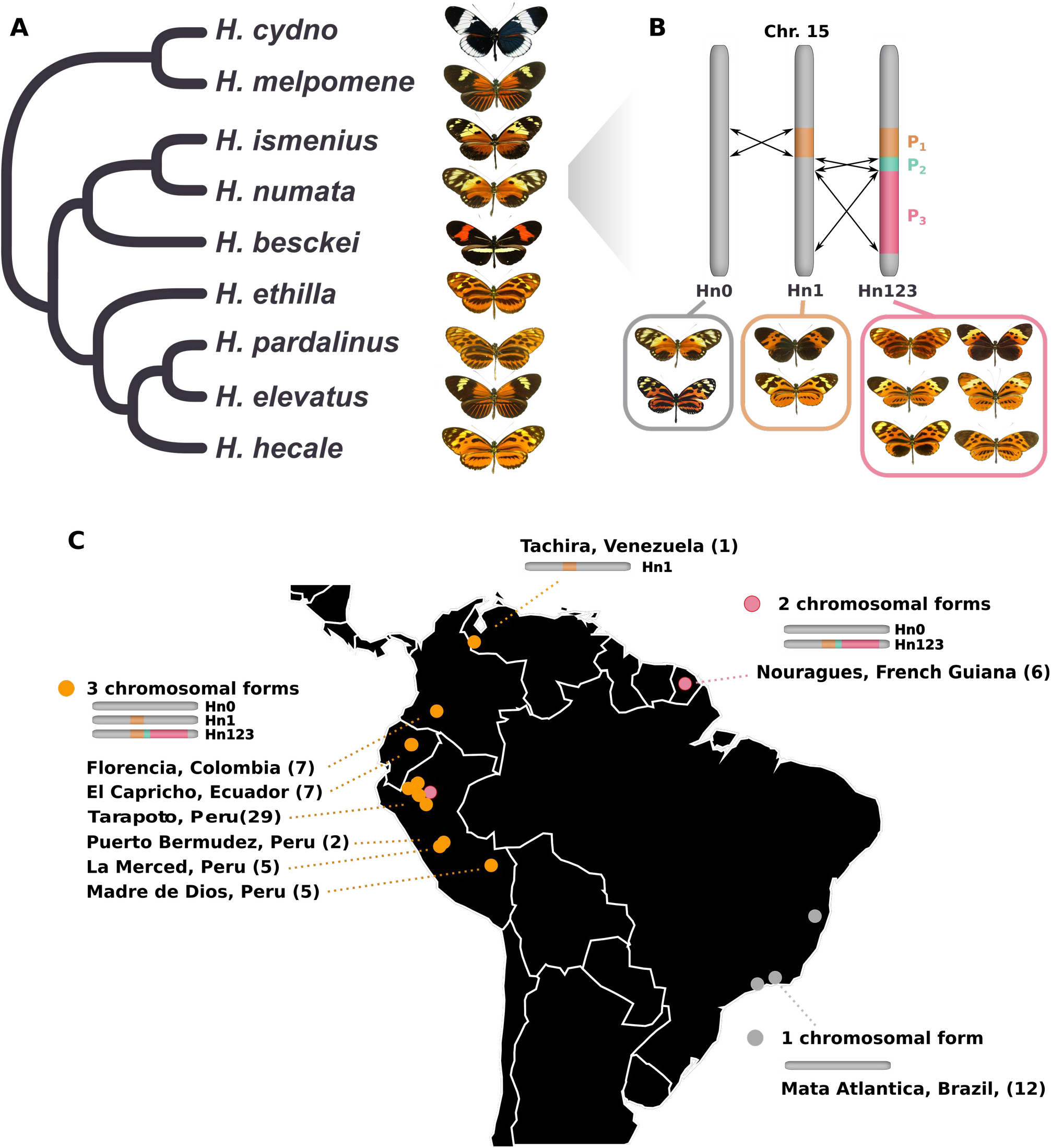
Genetic and population structure at the P supergene. **A**. Schematic phylogeny of the sampled species. It includes all members of the silvaniform clade and two outgroups, *H. melpomene* and *H. cydno*. **B**. Schematic description of the genetic structure of the P supergene. Three chromosomal arrangements coexist in *H. numata* and are associated with different morphs. **C**. Origin of *H. numata* specimens used for analyses and distribution of chromosome arrangements across the neotropics. Numbers in brackets indicate sampled specimens in each locality (the Tarapoto population lumps several neighbouring subsamples on the map)

*Heliconius numata* is widespread in the lowland and foothill tropical forests of the Amazon basin, the Guianas, and the Brazilian Atlantic Forest (Mata Atlântica), but the frequencies of the three chromosome arrangements vary across the range. Ancestral type Hn0 is fixed in the Atlantic Forest populations of Brazil (forms *robigus* or *ethra)*, but segregates at intermediate frequencies in all other *H. numata* populations throughout the range (forms *silvana* and *laura*) (**Fig 1C**). Chromosome type Hn1 is associated with the Andean mimetic form *bicoloratus* and is found in the Eastern Andean foothills of Ecuador, Peru, and Bolivia. Chromosome type Hn123 is associated with a large diversity of wing-pattern forms of intermediate allelic dominance, including *tarapotensis, arcuella* and *aurora*, and is reported from Andean, lowland Amazonian and Guianese populations. Inversion polymorphism is therefore structured across the range, with populations being fixed for the ancestral chromosome (Atlantic Forest, see Text S1 & Table S1-2), or displaying a polymorphism with two (Amazon-Guiana) or three (Andes) chromosomal types in coexistence (Joron et al. 2011). Monomorphic populations of the Atlantic Forest, devoid of rearrangements at the supergene locus, might represent the ancestral state displayed by *H. numata* populations before the evolution of the supergene via introgression (**Fig 1C)**.

The wing patterns of *H. numata* are subject to selection on their resemblance to local co-mimics (Chouteau et al. 2016), but the polymorphism is maintained by balancing selection on the chromosome types. Balancing selection is indeed mediated by disassortative mating favouring mixed-form mating (Chouteau et al. 2017) and is likely to have evolved in response to the deleterious mutational load carried by inversions, which causes heterozygous advantage in *H. numata* (Jay et al. 2021, Faria et al. 2019, Maisonneuve et al. 2019). The introgression of P_1_ and the formation of a supergene were associated with a major shift in the selection regime (Jay et al. 2018). The mating system also changed during or after introgression. These events may therefore have profoundly affected the population biology of the recipient species, *H. numata*. We investigate here whether the adaptive introgression of a balanced inversion is associated with a signature in genetic diversity and geographic structure. In particular, we predict that genetic diversity should be higher in *H. numata* than in closely related taxa. Similarly, nucleotide diversity should be higher in all polymorphic populations carrying either one segment (Hn1) or two (Hn1,Hn123) compared to the population that is monomorphic and only carries the non-inverted segment (Hn0) in the Brazilian Atlantic Forest. We analyse changes in the demographic history of the clade containing *H. numata* and closely related taxa, as well as their current patterns of diversity and demography, using three well separated populations of *H. numata* representing different states of inversion polymorphism. Our results suggest that following supergene formation, a change in the selection regime and mating system may have facilitated gene flow among morphs and had key consequences in current patterns of genetic structure.

## Material and Methods

We used here whole genome resequencing from 137 specimens of *Heliconius*, including 68 *H. numata*. Sampling included specimens from populations in the Andean foothills (3 chromosome types), from the upper Amazon (2 chromosome types), from French Guiana (2 chromosome types) and from the Brazilian Atlantic Forests (1 chromosome type) (Fig 1C; Table S3). Related taxa were represented by the sister species *H. ismenius*, found west of the Andes (parapatric to *H. numata*), by Amazonian representatives of the lineage *H. pardalinus* (donor of the inversion), *H. elevatus, H. ethilla, H. besckei* as well as *H. hecale*, and by *H. melpomene* and *H. cydno* as outgroups. Only Andean, Amazonian and Guianese populations of *H. numata* display chromosomal polymorphism, all other taxa being fixed for the standard gene arrangement (Hn0), or for the inverted arrangement Hn1 (*H. pardalinus*) (Jay et al. 2018). Hereafter, *H. numata* populations from the Andes, Amazon and French Guiana will be collectively referred to as “Amazonian”, and populations from the Atlantic Forest as “Atlantic”. Butterfly bodies were preserved in NaCl saturated DMSO solution at 20°C and DNA was extracted using QIAGEN DNeasy blood and tissue kits according to the manufacturer’s instructions with RNase treatment. Illumina Truseq paired-end whole genome libraries were prepared and 2×100bp reads were sequenced on the Illumina HiSeq 2000 platform.

Reads were mapped to the *H. melpomene* Hmel2 reference genome (Davey et al., 2016) using Stampy (version 1.0.28; Lunter and Goodson, 2011) with default settings except for the substitution rate, which was set to 0.05 to allow for the expected divergence from the reference of individuals in the so-called silvaniform clade (*H. numata, H. pardalinus, H. elevatus, H. hecale, H. ismenius, H. besckei* and *H. ethilla*). For *H. melpomene* and *H. cydno* belonging to the so-called *melpomene* clade, their genomes were mapped with a substitution rate of 0.02. Alignment file manipulations were performed using SAMtools v0.1.3 (Li et al. 2009). After mapping, duplicate reads were excluded using the *MarkDuplicates* tool in Picard (v1.1125; http://broadinstitute.github.io/picard) and local indel realignment using IndelRealigner was performed with GATK (v3.5; DePristo et al. 2011). Invariant and polymorphic sites were called with GATK HaplotypeCaller, with options --min_base_quality_score 25 --min_mapping_quality_score 25 -stand_emit_conf 20 --heterozygosity 0.015.

F_ST_, d_XY_ and π were calculated in overlapping windows of 25 kb based on linkage disequilibrium decay (*Heliconius* Genome Consortium 2012) using custom scripts provided by Simon H. Martin (https://github.com/simonhmartin), and the genome-wide average was calculated using our own scripts (available from https://github.com/angelesdecara). Distance in km between sampling sites was measured along a straight line, not taking into account potential physical barriers. Following Rousset (1997), in a 2-dimensional habitat, under a model of isolation by distance (IBD) differentiation, measured as F_ST_/(1-F_ST_), should increase as a function of the logarithm of distance. Therefore, we tested for the existence and intensity of an IBD signal among species and between populations of *H. numata* using a linear model. If IBD is stronger in species not polymorphic for the inversion we should observed significantly steeper slopes in these species. To test this, we measured IBD (*i*) within populations of each species separately, (*ii*) for all *H. numata* within the Amazonian region (excluding Atlantic forest populations) and (*iii*) for all *H. numata* including the Atlantic region. The slopes of F_ST_/1-F_ST_ versus log(distance) were calculated using the R package *lsmeans* (Lenth 2016); the slope difference among species or between populations within species was estimated with an ANOVA and its significance evaluated with function pairs of this package (Text S1 and see example script on github.com/angelesdecara/).

Admixture (Alexander et al. 2009) analyses were run on a subset of the 68 *H. numata* genomes, keeping only 15 individuals from Peru to have a more balanced representation of individuals across the geographic distribution. Filters were applied to keep biallelic sites with minimum mean depth of 8, maximum mean depth of 200 and at most 50% genotypes missing. We only kept 1 SNP per kilobase to remove linked variants using the thinning function in vcftools, and we obtained the optimal number of clusters using cross-validation for values of K from 1 to 10 (Alexander et Lange, 2011). Principal component analyses (PCA) were performed with the same filters as for admixture, using the same *H. numata* genomes as for the admixture analyses, using plink2 (Chang et al. 2015).

In order to estimate demographic parameters independently of the effect of selection on diversity, we performed stringent filtering on the dataset. We removed all predicted genes and their 10,000 base-pair flanking regions, before performing G-PhoCS (Gronau et al. 2011) analyses as detailed below. Repetitive regions were masked using RepeatMasker and Tandem Repeat Finder (Benson 1999). GC islands detected with CpGcluster.pl with parameters 50 and 1E-5 (Hackenberg et al., 2006) were also masked. Scaffolds carrying the supergene rearrangements (Hmel215006 to Hmel215028) were excluded, as were scaffolds from the sex chromosome (Z) and mtDNA, since those are expected to show unusual patterns of diversity due to selection and different effective population sizes.

We analysed the demographic history of *H. cydno, H. numata, H. ismenius, H. pardalinus* and *H. elevatus* with G-PhoCS, which allows for the joint inference of divergence time, effective population sizes and gene flow. In order to detect differences in demography correlating with the presence of the supergene in *H. numata*, we conducted analyses separating the Atlantic population of *H. numata* from Amazonian populations. G-PhoCS is an inference method based on a full coalescent isolation-with-migration model. Inferences are conditioned on a given population phylogeny (based on Kozak et al. 2015) with migration bands (i.e. priors in the migration rates) that describe allowed scenarios of post-divergence gene flow. The model assumes distinct migration rate parameters associated with each pair of populations, and allows for asymmetric gene flow. Given the computational burden of G-PhoCS, we selected two individuals per taxon or population, retaining those with the highest sequencing depth (see Table S3), taking into account their relative abundance. The input dataset consisted of 4092 genomic regions, each 1kb in length and spaced at approximately 30kb intervals (above the value at which LD decays at more than half of its value) and with genotypes in at least one of the two samples of each taxon We used as priors for coalescence times (τ) and genetic diversity (θ), Gamma functions with α=1 and β=100, and for migration rates α=0.002 and β=0.00001. These priors were chosen to allow good convergence while also ensuring non informativity. In order to calculate the highest posterior density interval, we used the library HDInterval in R, and to integrate such posterior densities we used the library sfsmisc in R. We rescaled the results using a mutation rate of 1.9e-9 µ/bp/generation (Martin et al. 2016) and 4 generations per year (i.e., g=0.25). Migration bands were considered significant following the criteria of Freedman et al. (2012): if the 95% HPD interval did not include 0 or if the total migration rate (i.e. migration rate times the duration of the migration band) was larger than 0.03 with posterior probability larger than 0.5.

### Demographic Reconstruction of population size changes, split and mixtures

G-phocs provides useful information across all species but i) does not allow to quantify the time scale of population size change, ii) is limited in the number of individuals it can handle and iii) displayed limited accuracy to distinguish Ne and *m* in a simulation study (Gronau et al. 2011). We thus constructed additional models to test the hypothesis that *H. numata* populations with inversion polymorphism display an increased effective population size due to disassortative mating. To test this, we used ∂a∂i (Gutenkunst et al. 2009) to reconstruct the demographic history of *H. numata* individuals from the Amazonian forest, quantify their historical changes in effective population size and test their divergence history from 1) *H. numata* from the Brazilian area, which do not carry the inversion; and 2) *H. pardalinus* individuals. We allowed for change in effective population size in both the ancestral population and daugther populations. The change in effective population size in *H. numata* associated with the change in mating system should be more recent than the time of introgression of the inversion into *H. numata*. To verify this hypothesis, we allowed for change in *Ne* of the daughter population at any time after the split. We tested different models of divergence with and without (asymmetric) migration and included the effect of linked selection and barriers to gene flow (Roux et al. 2016).

Since the conditions of historical divergence are not known, we tested a model of divergence with ongoing migration (IM) a model of divergence with ancient migration if gene-flow has stopped recently (AM) and, in the case of divergence into multiple refugia, a model of secondary contact (SC). We also included a model of Strict Isolation (SI) as a null model. The models shared the following parameters: the ancestral population of size *N*_*anc*_ can grow or shrink to a size N_*anc2*_ between T_*anc*_ and up until its splits at time *T*_*split*_ into two daughter populations of size *N*_1_ and *N*_2_. Under the SI model, no gene flow occurs between the two populations. Under AM, gene flow occurrs between T_*split*_ and T_*am*_ and is followed by a period of strict isolation. Under IM, gene flow occurs at a constant rate at each generation between the two populations. Gene flow can be asymmetric, so that two independent migration rates *m*_12_ (from population 2 to 1) and *m*_21_ (from population 1 to 2) were modeled. Under the SC model, the population evolved in strict isolation between *T*_*split*_ and until *T*_sc_ where a secondary contact occurs continuously up to present time. Gene flow is modeled as M = 2*N*_REF *_ *m*. Heterogeneity in migration rate was used to account for the accumulation of barriers to gene flow by defining two categories of loci, with varying migration rate. Those with a “neutral” rate of migration, and those in proportion P with a reduced rate of migration due to barriers. Similarly, heterogeneity in effective population size was used to account for linked selection by defining two categories of loci with varying effective population sizes (proportion 1-Q of loci with a “neutral Ne” and a proportion Q of loci with a reduced effective population size due to either selection at linked site). To quantify how linked selection effects reduced *N*_*e*_, we used a Hill-Robertson scaling factor (Hrf) to relate the effective population size of loci influenced by selection (Nr = Hrf * *N*_*e*_) to that of neutral loci (*N*_*e*_). A hierarchical approach was used to avoid over-fitting: first, we compared models assuming constant effective population size. Second, the best identified models were modified to incorporate population expansion or decline, as expected given the observed distribution of genetic diversity. Population expansion was implemented using two additional parameters for population 1 and population 2, allowing each population to either grow or decline exponentially at any time after their split from the ancestral population (controlled by parameters s1 and s2 for population 1 and 2 respectively) which defined the new time parameters (Tp1 and Tp2), which indicate the time of exponential change).

Models were fitted using the diffusion theory implemented in ∂a∂i which uses the SFS as a summary of the data. For a given demographic model, the SFS is computed using diffusion approximation and compared to the empirical SFS using AIC.

We used stringent filtering (GQ>30, 4 < mean depth < 80) and no missing data to keep high quality sites and remove potential paralogs or PCR duplicates exhibiting excessive read depth. To minimize linkage we subset our data to keep one SNP every 5kb. No MAF filter was used and singletons were kept to avoid ascertainment bias in estimates of demographic parameters. For each model, 32 independent replicate runs were performed and for each model, the run with the lowest AIC and ΔAIC was kept.

### Forward Simulations

In order to better understand the nature of the processes that generate higher genetic diversity in *H. numata* compared to closely related taxa, we used simulations to test the hypothesis that disassortative mating generates an increase in levels of genetic diversity at a genome-wide scale. We hypothesized that such level of genetic diversity is higher than expected under i) random mating (a model similar to panmixia) or ii) assortative mating, as observed between divergently-coloured geographic variants (“races”) in *H. melpomene* (Jiggins et al. 2004) or in *H. timareta* (Sanchez et al 2015). Few polymorphic populations have been studied in *Heliconius*, besides *H. numata*, but partial assortative mating based on colour differences was found in *H. cydno alithea* (Chamberlain et al. 2009).

To test our hypothesis we ran forward simulations under two models: one model with disassortative and one model with assortative matting using slim v3.6 (Messer et al. 2013). Both models were run using different strengths of mate choice, notably in order to compare the level of genetic diversity under random mating to the level observed under scenarios with strong mate choice. We attempted to choose both biologically and computationally realistic parameters for our models.

We simulated a stepping stone model with 10 demes in order to reflect the significant isolation by distance pattern observed here. Each deme was composed of 1,000 diploid individuals and connected by a (symmetric) migration parameter (m). Each individual received neutral and deleterious (ratio 16:6) mutations at a rate µ = 1e-8 µ/bp/generation (rescaled to µ = 1e-6 for faster simulation of a larger population). We simulated an individual with a pair of 1Mb chromosomes, including a single locus with 5 possible alleles with perfect dominance (allele 1 > allele 2 > allele 3 > allele 4 > allele 5) given 5 possible alternative phenotypes (referred hereafter as “morph”) based on patterns of dominance observed elsewhere (Le Poul et al. 2014). Each allele was fully linked (no recombination) with a given set of deleterious recessive mutations, generating overdominance at this locus so that polymorphism is always maintained. Local adaptation was introduced in the model through a single parameter defining randomly which morphs were favoured in each population. In each population, either 2 or 3 morphs benefited from a fitness advantage compared to the others. In *H. numata* up to 7 distinct wing-pattern morphs can co-occur within a single locality (Chouteau et al. 2017). Therefore, we allowed multiple morphs to coexist within a deme. Fitness varied between 0 (= null fitness for non-locally adapted individuals) and 1 (no reduction of fitness). We tested 4 possible values for this parameter (0, 0.25, 0.5 and 1).

Finally, disassortative mating was controlled by a mate choice parameter defining whether a morph would reproduce with another morph. The strength of the parameter varied between 0 (= complete disassortative mating, meaning that a given individual mates only with a different morph) and 1 (= no mating weight, meaning random mating). We tested 4 possible values for this parameter (0, 0.25, 0.5 and 1 (i.e. meaning random mating).

We run the model for 80,000 generations to reach demographic equilibrium and assess levels of synonymous diversity (π_s_). We tested all combinations of the 4 values for levels of disassortative mating and local adaptation and ran 10 replicates per combination in order to estimate the variance around π_s._

Similarly, we run a model with strict assortative mating, controlled by a parameter defining whether similar morphs reproduced together. The strength of the parameter varied between 0 (complete assortative mating where a given individual mate only with an identical morph) and 1 (where individual mates randomly with regards to the morph). We tested 4 possible values for this parameter (0, 0.25, 0.5 and 1). As for disassortative mating, all combinations of assortative mating and local adaptation values were tested. For each model we tested 3 values for the migration rate, m = 1e-4, 1e-6 and 1e-8, resulting in a total of 54 comparisons.

For graphical display in Figure 4, the values of assortative/disassortative mating were rescaled on a scale between -1 (complete assortative mating and 1 (complete disassortative mating) with 0 indicating random mating. Values between -1 and -0.25 indicated cases of assortative mating, while values between 0.25 and 1 represent disassortative mating.

## Results

Using cross validation error as a measure of the optimal number of clusters with Admixture, we found that K=2 was the optimal cluster number describing within-species genetic variation in *H. numata* (**Fig 2A**). One cluster corresponds to the Atlantic population, forming a well-differentiated genetic entity compared to all other *H. numata* populations. All Amazonian populations of *H. numata* showed remarkable uniformity. This pattern is consistent with the population structure inferred using microsatellite markers (Fig S1). Population structure revealed by PCA is in line with the admixture analysis (**Fig 2B**). Individuals from the Atlantic populations of *H. numata* clustered together to one side of the first PCA axis, whereas all other individuals from all other populations clustered to the other side. The second axis of the PCA separates individuals from French Guiana from the other samples of the upper Amazon. This separation was not found with Admixture (i.e. with K=3) from the complete dataset. To better investigate the existence of a hierarchical population structure, we excluded individuals from the Atlantic populations and compared individuals from French Guiana to a randomly sampled set of Peruvian individuals. In this case we found a clear separation in two groups corresponding to French Guiana and Peru (Fig S2A). The same pattern was observed when replacing Peru by Colombia or Ecuador (Fig S2B,C). In accordance, pairwise genome-wide estimates of differentiation (*F*_*ST*_) between *H. numata* populations showed elevated values when comparing the Atlantic population to other populations, low values between French Guiana and other Amazonian populations, and were lower when comparing pairs of Amazonian populations outside of French Guiana (**Fig 2C**, Table S4). For instance, the population from La Merced in Peru shows an *F*_*ST*_ = 0.032 with the population from French Guiana at a distance of 3019km, but an *F*_*ST*_ = 0.311 (an order of magnitude higher) with the Atlantic population at a similar distance. The comparison between La Merced and Ecuador was even lower (*F*_*ST*_ = 0.0159). Isolation by distance among Amazonian populations of *H. numata*, estimated using the proxy *F*_ST_/1-*F*_*ST*_ ∼ log10(km) was significant (R^2^ = 0.41, p = 1.61e-06, slope = 0.02). Comparison among other species did not reveal any significant IBD (R^2^ = 0.01, p = 0.29, slope = 0.12). An analysis of the slope revealed a lower rate of increase in F_ST_ with distance in *H. numata* compared to all other taxa (**Fig 2C**, Table S4, Supp Text S1). By contrast, IBD between Atlantic and Amazonian populations of *H. numata* is close to what is observed in other species, and not significantly different (see Supp. Text S1).

**Figure 2.**
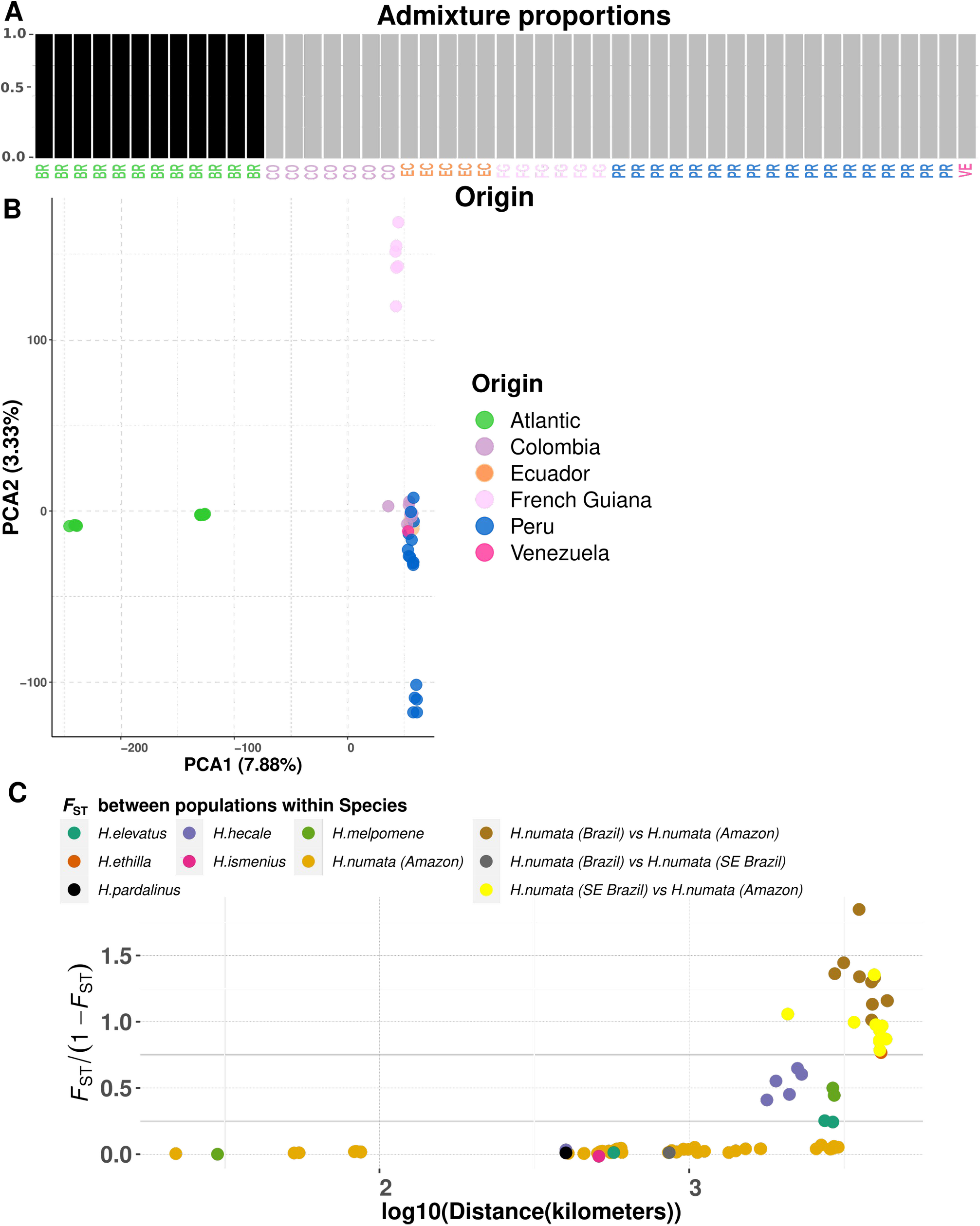
*H. numata* is characterised by low population structure. A. Admixture plot for *H. numata*. The optimal cluster number for *H. numata* is two, and it splits *H. numata* into two categories, whereas they come from Atlantic forest or the Amazon. BR=Brazil (Atlantic), PR=Peru, VE=Venezuela, CO=Colombia, EC=Ecuador, FG=French Guiana. B. Principal component analysis computed on whole genome SNP. Color code match those given in the admixture label of panel A. C. Relationship between genetic differentiation (Fst/(1-Fst)) and logarithm of geographical distance. Fst is measured between morphs/populations of the same species. *H. numata* populations from the Amazon show low isolation by distance when compared to related species.

Analyses of genetic diversity show that all populations of *H. numata*, except those from the Atlantic Forest, have a similarly high genetic diversity (**Fig 3A**). By contrast, closely related *Heliconius* taxa show lower genetic diversity (**Fig 3A**). These patterns are similar to those obtained using G-PhoCS to analyse the demographic histories in a phylogenetic context, where Amazonian populations of *H. numata* show higher population sizes compared to Atlantic Forest populations (**Fig 3B**, Table S5). G-PhoCS analyses also show a demographic history in which gene flow plays a crucial role (Table S6). For instance, our analyses show significant gene flow right at the beginning of the divergence between *H. ismenius* and the other silvaniforms, as well as in the divergence between *H. pardalinus* and *H. elevatus*. The effective population sizes inferred from Atlantic genomes are one order of magnitude lower than those obtained using *H. numata* populations from other localities (**Fig 3A** and Table S5). In our cladogram, the increase in *H. numata* population size is restricted to the Amazonian branch, excluding Atlantic populations.

**Figure 3.**
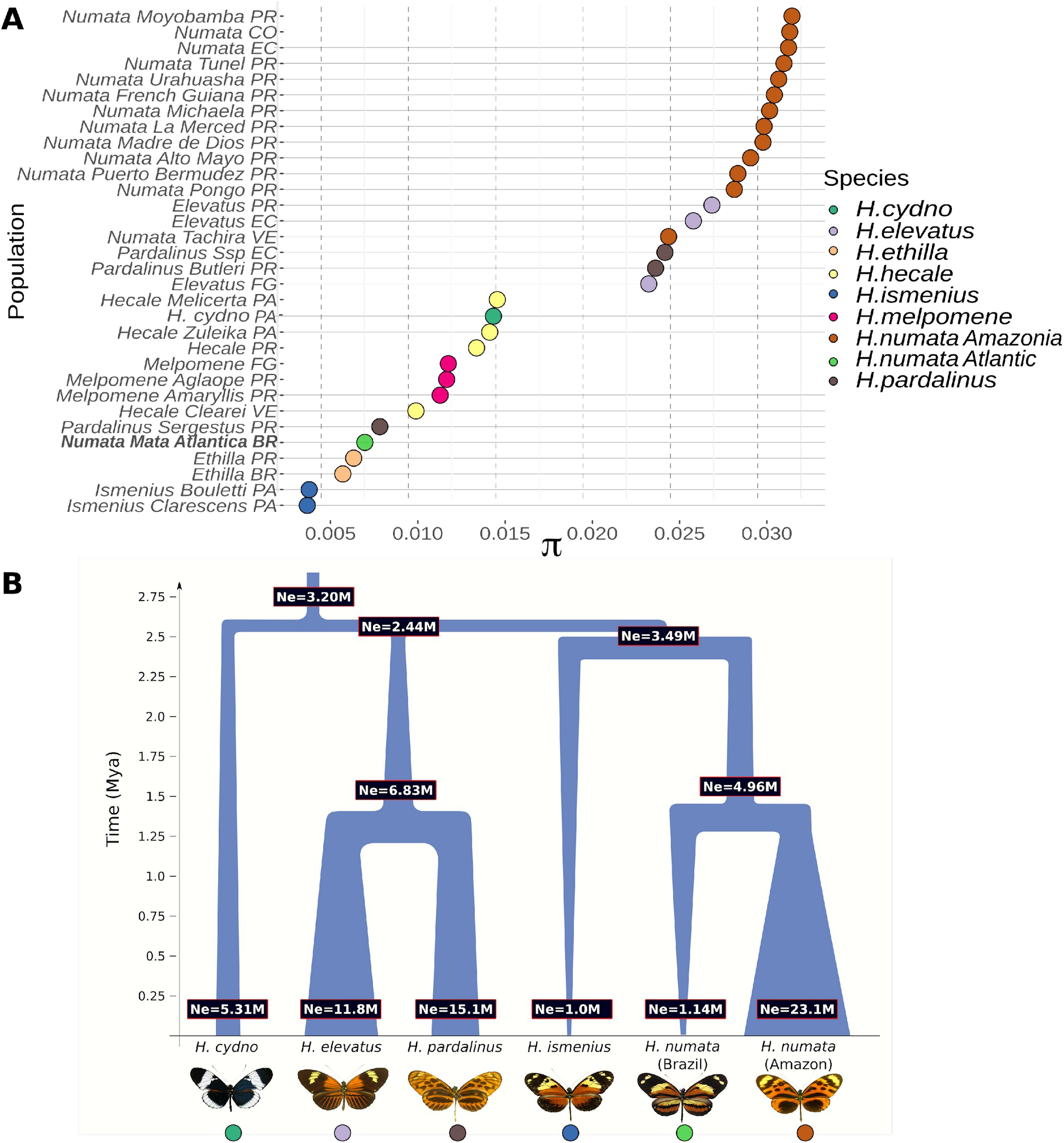
Variation in present and past effective population size in *Heliconius* species. **A**. Variation in Pi in several *Heliconius* populations, showing higher genetic diversity in *H. numata* populations from the Amazon than other taxa. Population names indicates their origin as in Figure 2 (e.g. PR=Peru), with the addition of PA=Panama. The *H. numata* population with a lowest diversity is the one from the Atlantic forest (Brazil). **B**. Schematic representation of Gphocs results (presented in table S5-6). Gene flow was modelled but not represented graphically for clarity. showing that Amazonian populations of *H. numata*, which have the P supergene, show a dramatic increase in population size posterior to their split with the Atlantic populations of Brazil, which lack the supergene.

**Figure 4.**
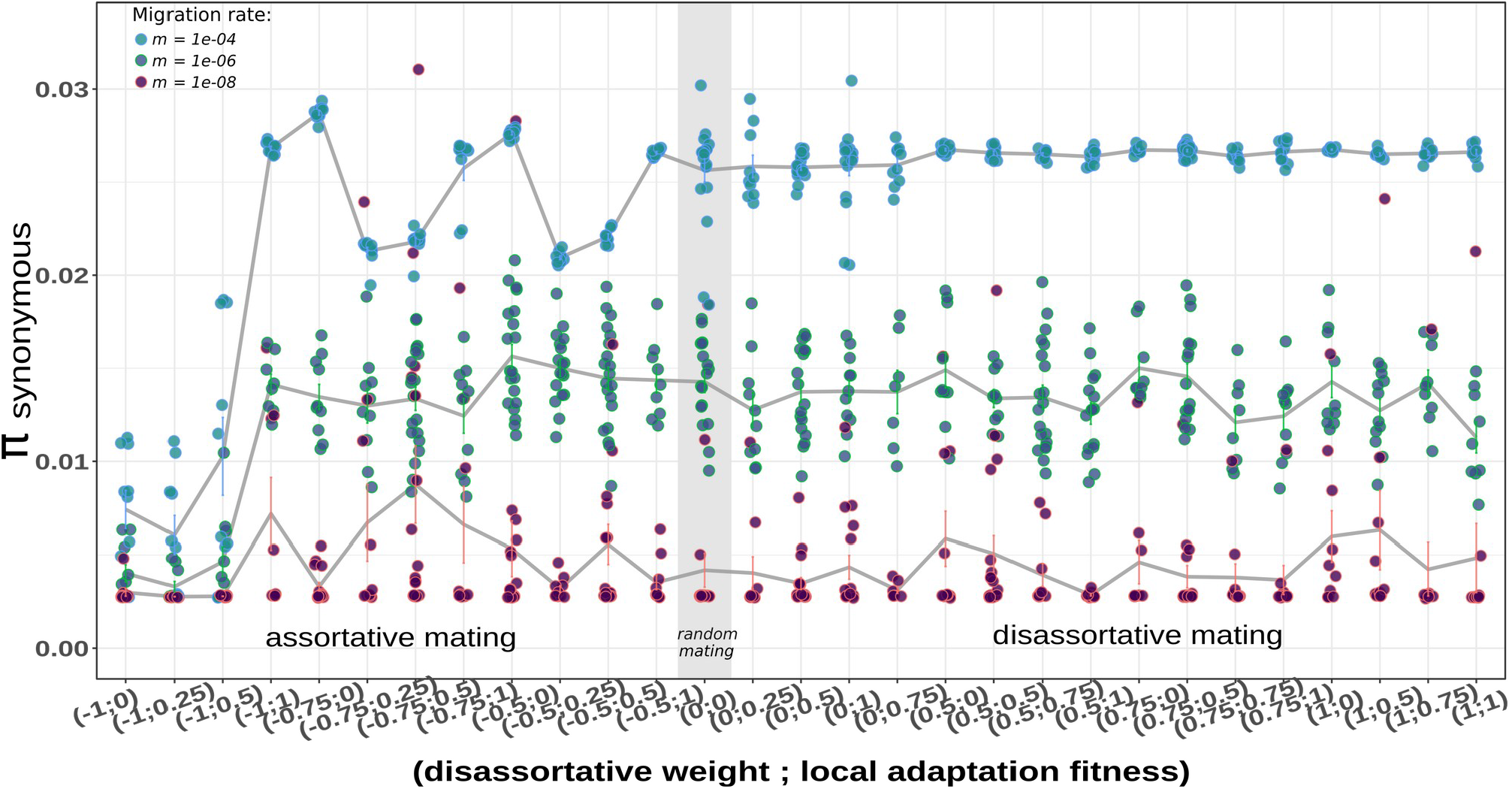
Weak but significant differences in synonymous nucleotide diversity (π_S_) emerged at a genome-wide scale under divergent selection and mating regime. Results from forward simulations of 10 replicate per parameter combination. A set of 10 demes with varying levels of migration, local adaptation fitness and different mating strategy (from assortative mating to disassortative mating) are presented. Shown are levels of synonymous diversity obtained under each combination of parameters. Parameters on the left part of the brackets display the disassortative mating weight (from -1 to 1). Parameter on the right side of the brackets displays the fitness value for local adaptation. A left value of -1 in the bracket means complete assortative mating and 0 means random mating. 1 = complete disassortative mating. A right value of 0 in the bracket means a fitness of 0 for non locally adapted individuals in a deme. A value of 0.5 means a reduced fitness of 0.5 relative to the maximum value. A value of 1 means no loss of fitness.

**Table 1:**
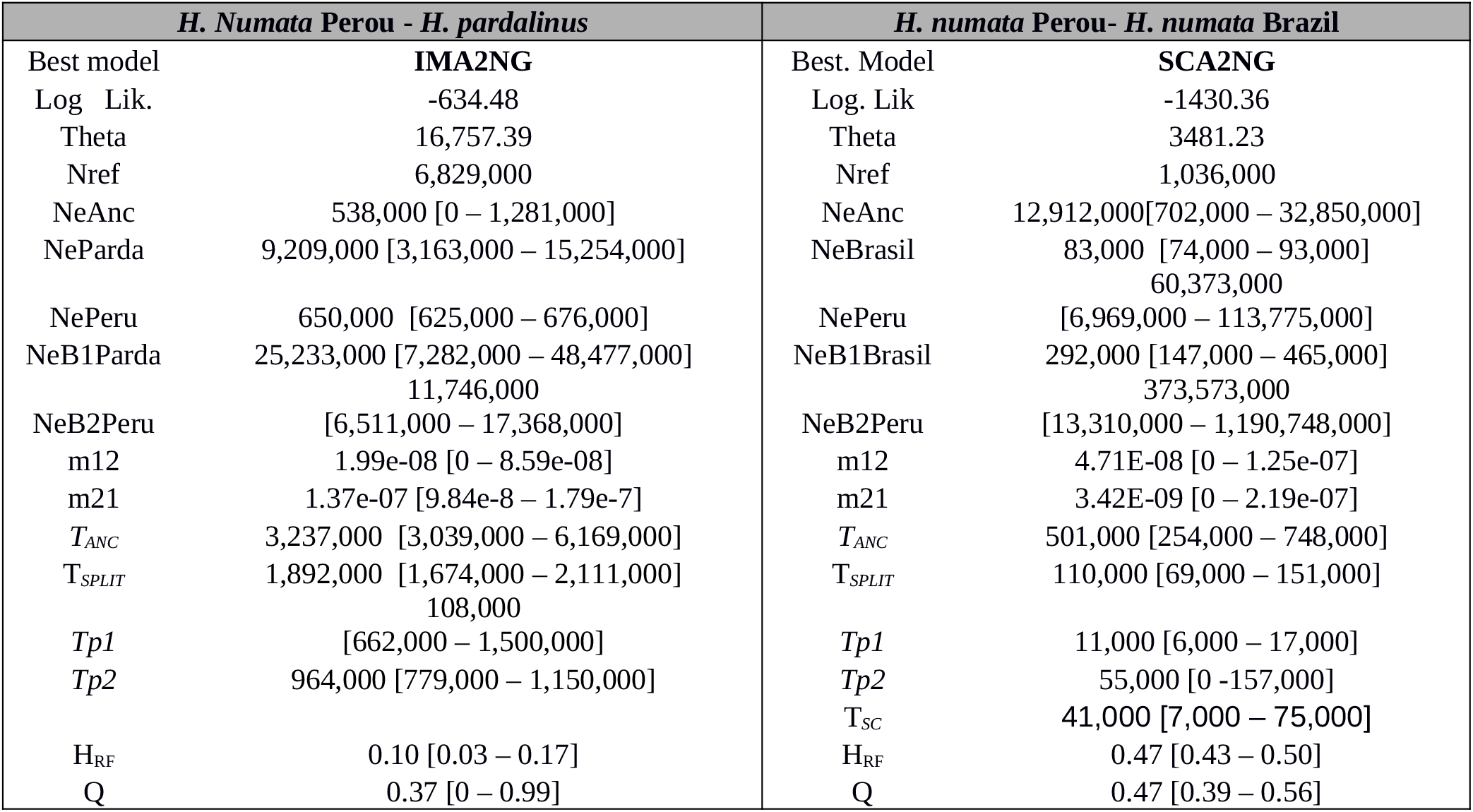
Estimates of demographic parameters for each best model in each comparison. Biological parameters assumed a mutation rates of 2.9e-09 µ/bp/generation. NeAnc, = Effective population sizes for the ancestral population. NeParda, NePeru and NeBrazil corresponds to effective population size of *H. pardalinus H. numata* in Peru and *H. numata* in Brazil respectively. NeB1Parda and NeB2Peru, NeB1Brasil provides effective population size for the population *H. pardalinus H. numata* in Peru and *H. numata* in Brazil respectively after their expansion. Hrf = Hill-Robertson factor reflecting the reduction in effective population size due to linked selection. Q = proportion of the genome with a reduced effective population size. m12 = migration from population 2 into population 1. m21 = migration from population 1 into population 2 (scaled according to 2N*ref*m_ij_). T_ANC =_ Time of population size change in the ancestral population, Tp1 = Time of size change in population 1 and Tp2 = Time of size change in population 2. T_ANC_, T_SPLIT_and Tsc, Tp1, Tp2 are provided assuming four generations per year.

### Demographic reconstruction from ∂a∂i

The model selection procedure based on AIC gave higher support for a model of secondary contact (SC) in the pairwise comparison between *H. numata* from Peru and *H. numata* from Brazil. The pairwise comparison between *H. numata* and *H. pardalinus* supported a model of divergence with continuous gene-flow (IM) (Table S7, Fig S3). All models supported an expansion occurring in the ancestral population, followed by further growth of the *H. numata* carrying the inversion supergene to reach a size of several millions, which was by far the largest effective size compared to all other species. This stands in stark contrast with the results observed in the samples from Brazil (which do not harbor the inversion) (Table 2). Accordingly, *H. numata* populations from the Atlantic forests of Brazil appear to have been subject to a bottleneck at the start of their divergence from Amazonian populations, followed by exponential growth, suggesting a strong (and recent) founding event, leading to a comparatively smaller population size than that observed in the rest of *H. numata*. It is worth noting however that effective population size was hard to estimate in pairwise comparisons between *H. numata* from Peru and SE Brazil. Indeed, parameter uncertainty was large, and model residuals (Fig S3) were also large. Our results indicated that *H. pardalinus* displayed an initially large population size followed by a comparatively smaller size expansion than *H. numata* (Table 2). There was a large variance in estimates of effective population size in *H. numata* from Peru in the two inferences (with *H. pardalinus* or with Brazilian *H. numata*). Yet, in bothinferences, the effective population sizes of the Peruvian *H. numata* were large (greater than 11 million). Estimates of current effective population sizes are therefore qualitatively similar to those from G-phocs.

### Forward simulations

Forward simulations under different levels of local adaptation (controlled by the strength of divergent selection), disassortative mating and migration are displayed in **Fig 4B**. The same results under a model of assortative mating involving different levels of selection and migration are displayed in **Fig 4A**. Overall, synonymous genome-wide nucleotide diversity (π_S_) was higher in 73 % of the models including disassortative mating (average π_S_ = 0.0145) when compared to their equivalent under assortative mating (average π_S_ = 0.011), a weak but significant difference (p <0.01). In summary, modest differences were observed among models with different strength of divergent selection or disassortative mating, the most influential variable being the rate of migration (**Fig 4**).

## Discussion

Our results suggest that populations displaying inversion polymorphism in the *P* supergene in *H. numata* also display distinctive population demography and gene flow. Differences in demographic and differentiation regimes associated with structural variation at this locus are revealed when comparing polymorphic populations of *H. numata* to closely-related monomorphic taxa, such as (1) peripheral populations of *H. numata*, (2) sister taxa, and (3) inferred ancestral lineages. This suggests that the existence of a mimicry supergene controlling polymorphism in *H. numata* may be associated, in time and in space, with major differences in population biology. We hypothesize this to be due to a change in the balancing selection regime due to heterozygote advantage (Jay et al. 2021) and in the associated evolution of disassortative mating (Chouteau et al. 2017) following the onset of inversion polymorphism, causing direct effects on ecological parameters such as gene flow, immigration success and effective population size. Testing this hypothesis through forward simulation yielded mixed evidence for a genome-wide effect of this disassortative mating, especially when compared to a simple model of random mating.

Our analyses show large-scale variation in genetic diversity among closely related taxa in this clade of *Heliconius* butterflies. Within *H. numata*, genetic diversity found in polymorphic Amazonian populations is ∼4 times higher than in populations from the Atlantic Forest. Generally, Amazonian populations of *H. numata* harbour the highest genetic diversity in the entire *melpomene*/silvaniform clade, which contrasts with the low diversity found in the most closely related taxa such as *H. ismenius* or *H. besckei*. Inferring historical demography during the diversification of the *H. numata* lineage reveals that the large effective population size in that species is only associated with the branch representing polymorphic, Amazonian *H. numata* populations, while internal branches all show very low diversity estimates. This suggests that ancestral and putatively monomorphic populations of *H. numata* were similar in their diversity parameters to current sister species *H. ismenius* populations, or to current peripheral Atlantic *H. numata* populations. Although low-diversity lineages could have lost diversity due to recent events such as strong bottlenecks, as estimates of effective population size from ∂a∂i indicated for the Atlantic population. Nevertheless our ∂a∂i estimates do suggest that the Amazonian populations of *H. numata* underwent a dramatic increase in effective population size posterior to their split with Central American (*H. ismenius*), *H. pardalinus* and the Atlantic populations. Those findings are in agreement with G-Phocs analyses. The Amazonian branch of the *H. numata* radiation is characterized by the long-term maintenance of inversion polymorphism, triggered by the introgression of a chromosomal inversion about 2.2 Ma ago. Therefore, the major shift in demography between Amazonian and Atlantic populations indeed appears to coincide in time, at least in the broad sense, with the occurrence of inversion polymorphism, even though the lack of replication of this event impedes firmly establishing causality.

Another striking result is the lack of genetic structure displayed by *H. numata* across the Amazon, with all Amazonian populations forming a single genetic cluster. Only Atlantic Forest populations stand out and display high differentiation with other *H. numata* from the rest of the range. French Guiana and Peruvian populations, separated by over 3000 km across the Amazon, are weakly genetically differentiated compared to pairs of populations at comparable distances in other species, and show only modestly stronger differentiation than pairs of *H. numata* populations taken at short distances. *Heliconius numata* populations from the Amazon show significantly lower isolation by distance than all other taxa, as measured by the change in *F*_*ST*_ across distance (*F*_*ST*_/km) (Fig. 2C), with a very distinctive, flat slope of isolation by distance. The only exception is found when comparing Amazonian populations with Atlantic Forest populations of Brazil, displaying a level of differentiation in line with that of pairs of populations at similar distances within other taxa.

Effective population size is affected by census size, mating system, and the force and type of selection acting on traits (Charlesworth 2009). Selection is often viewed as a force only affecting the genetic variation around specific, functional loci in the genome, but it may also affect whole genome diversity, for instance when its action is sufficient to modify local demography or mating patterns. In *H. numata*, morphs and therefore inversion genotypes show disassortative mate preferences, i.e., they preferentially mate with individuals carrying different chromosome types (Chouteau et al. 2017). Disassortative mating enhances heterozygosity and the mating success of individuals expressing rare alleles (negative frequency dependence) (Knoppien 1985; Hedrick et al. 2018). Consequently, immigrants expressing rare, recessive alleles have a mating advantage in *H. numata*. Disassortative mating associated with the supergene should therefore bring an advantage to immigrant genomes in LD with recessive supergene alleles, possibly enhancing genome-wide gene flow. Supergenes are also characterised by single-locus Mendelian inheritance, by which mimicry phenotypes are maintained in the face of recombination, even after immigration. Effective migration regime in populations harbouring a mimicry supergene is therefore likely to be quite different to that observed in other mimetic taxa such as *Heliconius melpomene* or *H. erato*, in which mimicry variation is controlled by multiple loci with diverse dominance patterns. In those taxa, hybrid offspring display recombinant patterns breaking down mimicry, even after multiple generations of backcrossing, and pure forms mate assortatively with respect to wing pattern (McMillan et al. 1997, Mallet et al. 1998, Jiggins et al. 2001); both processes select against mimetic variants migrating from adjacent areas with distinct warning patterns. The expectation is that immigrant genomes should be consistently associated with mimicry breakdown in the case of multilocus architectures, which should translate into a reduction of the effective migration genome-wide, compared to situations with polymorphic mimicry supergenes. In *H. numata*, the evolution of a polymorphic mimicry supergene and disassortative mate preferences could therefore explain the relative lack, compared to other *Heliconius* taxa, of differentiation among polymorphic populations, even across large distances. Furthermore, enhanced gene flow could also cause an increase in effective population size estimates (Slatkin 1987), putatively explaining why polymorphic populations of *H. numata* harbour the highest genetic diversity, and display the highest *Ne* estimates in the entire *melpomene*-silvaniform clade of *Heliconius*. These hypotheses are also supported by our forward simulation which revealed an effect of the mating system. Yet, most of the difference in genetic diversity was due to a assortative mating reducing genetic diversity rather than a diversifying effect of disassortative mating. In addition, our results also suggest that a simple model of random mating may explain well the data, thus purely demographic expansions may also generate high genetic diversity and high effective population size, as observed from our ∂a∂i demographic modelling. Overall, the change in genetic diversity due to mating system variation is limited compared to the effect of migration. Indeed, our simulation results indicated that migration between demes has the stronger effect on synonymous diversity. In addition, the magnitude of difference in the simulation is weak relative to the observed differences of genetic diversity between *H. numata* from Brazil and *H. numata* from Peru. Whether these differences between our simulations and those empirical observations can be directly compared is not straightforward however given the simplified assumption required by our modelling approaches. In all cases, our demographic simulations with ∂a∂i, do suggest a strong effect of founder events/bottlenecks in reducing the genetic diversity of the Brazilian population. Another limit of our model is that we tested the effect on genetic diversity on a 1 Mb segment of a single chromosome, not on unlinked chromosomes. Although testing the effect on unlinked data would be relevant, it would substantially increase the compute time, so we choose to focus on this simpler model and leave the question of large scale effect for future investigations.

Alternative processes may also contribute to the observed patterns. Amazonian and Atlantic populations may differ in other aspects that could also result in differences in genetic diversity. Habitat availability and structure may be different, possibly entailing differences in the maintenance of diversity. The Atlantic Forest is vast in area, but may represent a smaller biome compared to the Amazon, and is isolated from the bulk of the range of *H. numata*, which could result in populations displaying characteristics of peripheral populations with smaller effective population sizes (Eckert et al. 2008). Reduced effective population size is supported by our data. One major caveat associated to our inference remains the small number of individuals (n = 12) from the Atlantic Forest. Genetic diversity might be underestimated, notably if populations have a history of fragmentation in this area. The other *Heliconius* species in the clade have much in common with *H. numata* in terms of habitat and general ecology, yet their niche and life-history specificities and their phylogenetic histories may result in consistent differences with the polymorphic *H. numata* populations. All those specificities may contribute to the observed pattern in which polymorphic Amazonian populations of *H. numata* display high effective population size and a weak geographic structure in genome-wide genetic variation. Yet this pattern of variation correlates parsimoniously with the evolution of a supergene causing disassortative mating and single-locus control of mimicry variation in Amazonian *H. numata* populations, which provides an elegant mechanism explaining their differences with extant and ancestral closely-related lineages. However, we cannot rule out a role for conjectural differences in ecology and geography with all other taxa.

In conclusion, our results show a remarkable contrast in the demography and differentiation of populations within the Amazonian range of *H. numata* compared to closely related taxa and ancestral lineages, as well as with other taxa in the *melpomene*/silvaniform clade. The distinctiveness of this widely polymorphic species in the clade is consistent with the hypothesis that the evolution of a supergene maintained by balancing selection represents a major transition in this lineage, triggering changes in genome-wide patterns of diversity and population ecology over the last 2 million years since its formation. If this hypothesis is correct, the evolution of a locus under balancing selection may therefore feed-back on population ecology and diversification, and consequently on speciation.

Eco-evolutionary feedbacks between changes in genomic architecture and the ecological parameters of populations are still not well understood and few cases have been studied. The evolution of self-incompatibility in plants, affecting the rules of mating and feeding back on population ecology, connectivity, and demography, may be one example, but effects of the evolution of trait genetic architecture on population ecology may be more common than previously thought. In our study, more work on the determinants of variation in effective population sizes in the genus *Heliconius* is needed to determine the precise impact of the supergene on demography in *H. numata*. We believe that our results emphasise a potential link between genomic architecture, selection and demography, and should inspire future theoretical and modelling studies. Overall, our result suggests that balancing selection maintains structural polymorphisms affecting life-history traits may have a profound influence on species ecology.

## Contributions

MARdC, PJ, QR and MJ designed the study and wrote the manuscript. BH, AVLF, TTT, RRR, KLSB provided the Atlantic samples. CS provided the Colombian samples. MARdC, PJ and QR performed genomic analyses and simulations with input from AW. MARdC, PJ, MJ, FPP and MC collected the Peruvian and Ecuatorian samples. MC performed microsatellite analyses and organized fieldworks and butterfly rearing. All authors contributed to editing the manuscript.

## Aknowledgements

This work was funded by grants Hybevol (ANR-12-JSV7-0005) and Supergene (ANR-18-CE02-0019-01) from the Agence Nationale de la Recherche and European Research Council Grant MimEvol (StG-243179). We acknowledge the Genotoul and the Montpellier Bioinformatics Biodiversity (MBB) platforms for providing us with calculation time. We thank Dr. Vitor Becker, at the Serra Bonita Reserve (Bahia), Alexandre Soares, at the MN/UFRJ (Rio de Janeiro) and Dr. Marcelo Duarte at the MZ/USP (Sao Paulo) for their contribution to the collection of butterflies in Brazil. Field collections in Colombia were conducted under permit no. 530 issued by the Autoridad Nacional de Licencias Ambientales (ANLA). Field collections in Brazil were conducted under ICMBio permit SISBIO-10438. We are grateful to Marianne Elias and Violaine Llaurens for comments and discussions. AVLF acknowledges support from Fundação de Amparo à Pesquisa do Estado de São Paulo – (FAPESP) (Biota-Fapesp grants 2011/50225-3, 2013/50297-0 and 2021/03868-8) and Conselho Nacional de Desenvolvimento Científico e Tecnológico (CNPq) (421248/2017-3 and 304291/2020-0). KLSB acknowledges the financial support of FAPESP Process # 2012/16266-7. Brazilian specimens are registered under SISGEN (A701768).

## Data, scripts, code, and supplementary information availability

The raw sequence data were deposited in NCBI SRA and accession numbers are indicated in Supplementary table 3. Scripts for ∂a∂i inferences are available at https://github.com/QuentinRougemont/DemographicInference

Scripts for forward simulation are available at https://github.com/QuentinRougemont/slim_heliconius Additional script are avaiable at https://github.com/angelesdecara

Filtered vcf and additional script to reproduce all figures are available at 10.5281/zenodo.7319912

## List of Supplementary Materials

Table S1-6

Fig S1-2

Text S1

