## supplementary Materials for "Balancing selection at a wing pattern locus is associated with major shifts in genome-wide patterns of diversity and gene flow"

### Supergene formation is associated with a major shift in genome-wide patterns of diversity in a butterfly

**This file contains:**

Figure S1 to S5

Table S1-S8 (Table S3 provided as a separate file)

Text S1

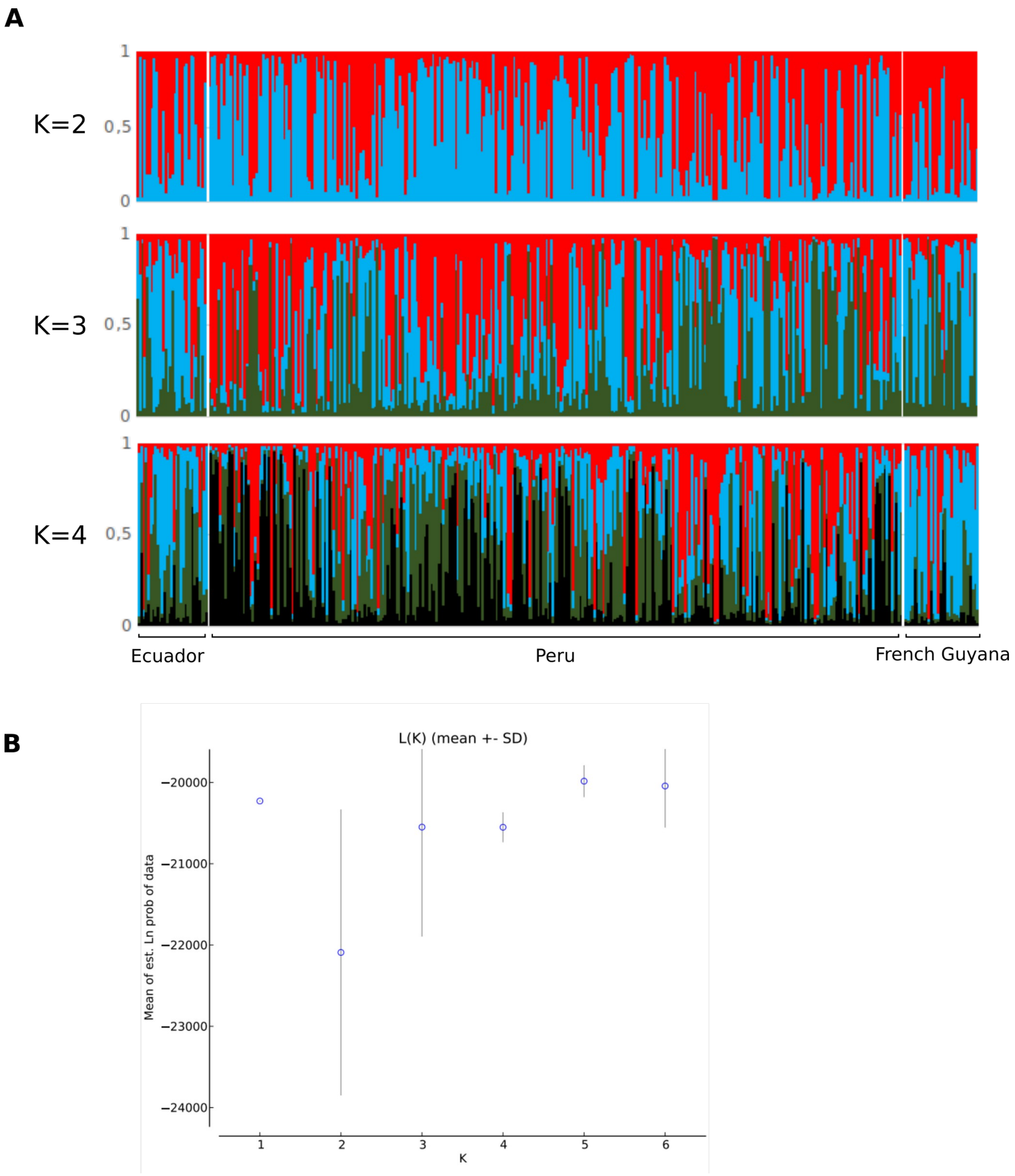


**Figure S1 | Genotype assignation plot using Structure showing general lack of population structure across the Amazon in *H. numata*.**

Detection of the number of groups (K) in STRUCTURE analysis of 360 *H. numata* individual genotyped at 14 microsatelites loci[1]. **A,** CLUMPACK[2] results following 20 STRUCTURE[3] run for each K ranging from 2 to 4 and setting both the length of burning period and the number of MCMC reps after burning at 100000. **B,** Mean value of log probability of the data L(K), as function of K ranging from 1 to 6 estimated by the program STRUCTURE HARVESTER[4] and following the method of Evano et al. (2005)[5].

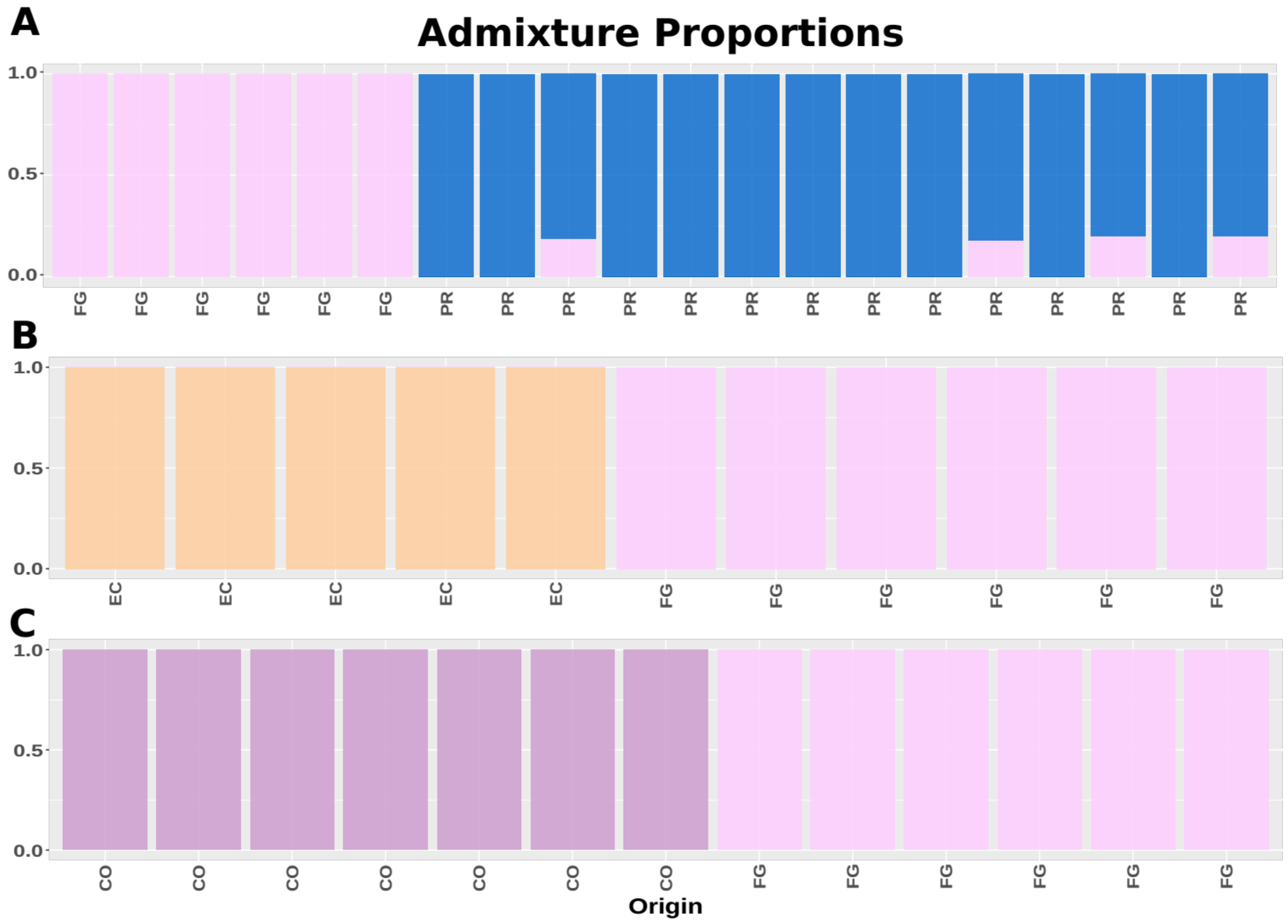


**Fig S2| Admixture analyses revealed substructure between French Guiana and other subgroups.**

Analysis was performed by sampling random individuals from Peru (PR) in panel A; or all individuals from other localities, namely Ecuador (EC) in panel B and Columbia (CO) in panel C.

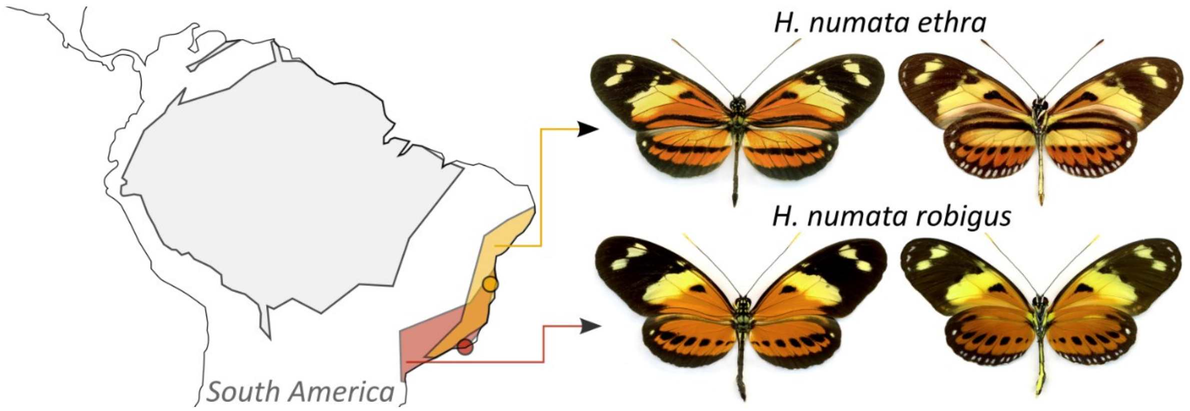


**Figure S3 | Geographic distribution, sampling sites and phenotype of the forms *H. numata ethra* (yellow area) and *H. numata robigus* (red area).**
Orange and red circles indicate sampling sites for *H. n. ethra* and *H. n. robigus*, respectively. The dorsal (left) and ventral (right) views of a specimen of each race are shown. The grey area on the map represents the range of distribution of the rest of the *H. numata* races. Geographic distribution of the races is according to Rosser *et al.* 2012.

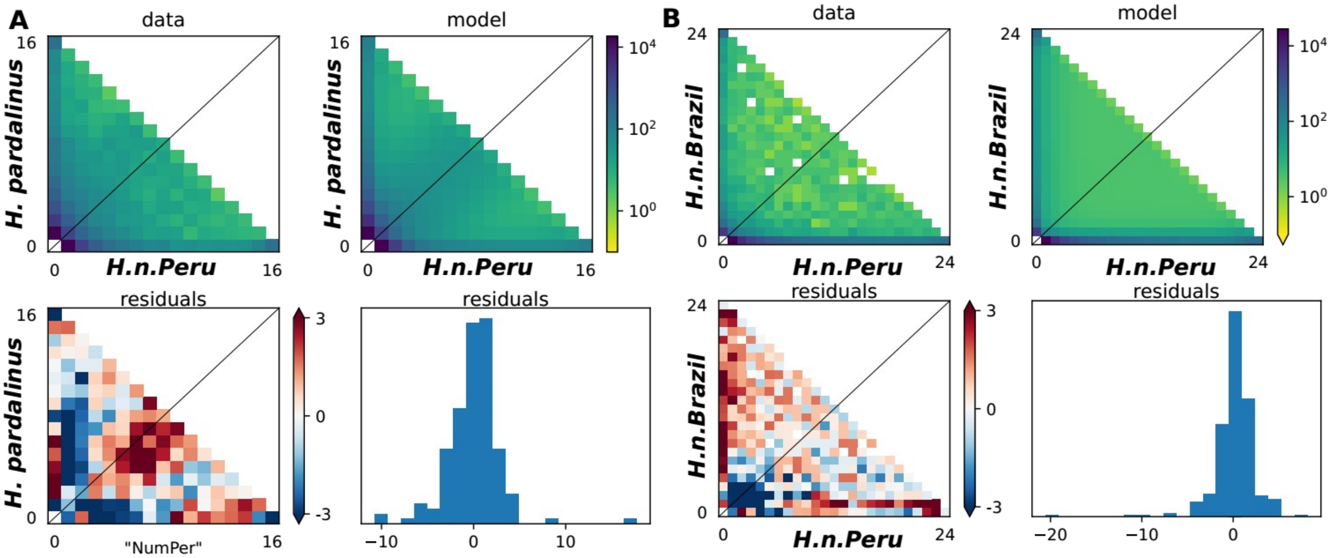


**Figure S4 | Differences between observed and predicted jSFS along with model residuals from dadi for the best models.** A) Comparison between H. *numata* from the Amazonian rain-forest in Peru and *H. pardalinus.* The best model is one of secondary contact with linked selection and population size change (SCA2NG). Migration was highly asymmetric. Site frequency spectrum for Peru in panel A) was down sampled to match the number of individuals in *H. pardalinus.*
B) Comparison between *H. numata* from Mata Atlântica in Brazil versus *H. numata* from the Amazonian forest in Peru. The best model is one of isolation with migration with linked selection and population size change (IMA2NG). B)

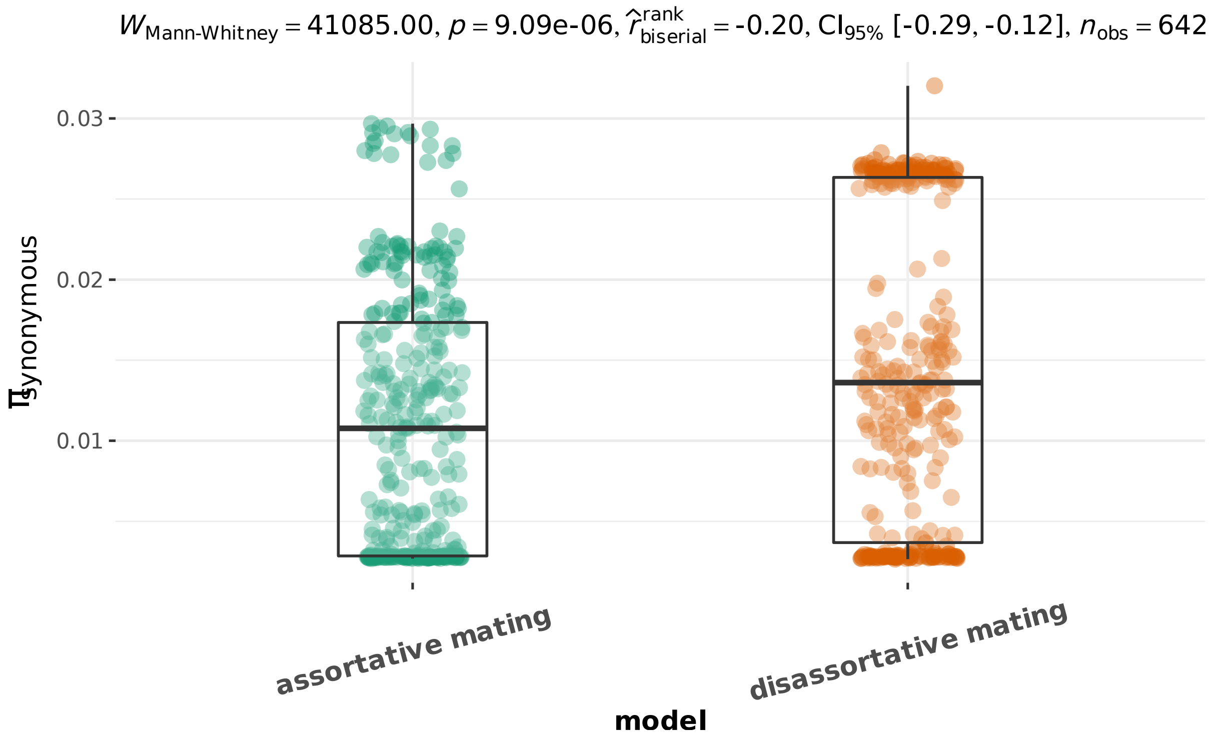


**Figure S5 |** Slightly higher synonymous diversity in models of disassortative mating compared to models of assortative mating (p <0.01). Most of the difference is due to models with m = 0.0001 as seen on Figure 4.

**Table S1. Band size of the *cortex* amplicons corresponding to the different gene orders at the supergene *P* in Peruvian populations of *H. numata***

| | | Expected band size (bp) | | |
|---|---|---|---|---|
| | | Arrangement *Hn0:* | Arrangement *Hn1* | Arrangement *Hn123* |
| First set of *cortex* primers (amplicon 1)[1] | F: CGCAACGTTATCGCCTAGATAGGTTCG<br>R: AANGCGAAASMACTGAYAACACGWG | ~500 | ~1100 | ~800 |
| Second set of *cortex* primers (amplicon 2)[2] | F: CGTAGCGACCCGAGATTCTT<br>R: AATACATGGCCACAGTTGATTC | ~380 | ~920 | ~650 |

[1]Jay et al. 2021 [2] Saenko et al., 2019

**Table S2**. Supergene alleles in the *numata* populations from the Brazilian Atlantic Forest

| **Form** | **Population** | **Specimen** | ***cortex* amplicon 1 (bp)** | ***cortex* amplicon 2 (bp)** | **Summary** |
|---|---|---|---|---|---|
| *H. numata ethra* | Reserva Serra Bonita | BH16-0020 | Null | Null | **Arrangement *Hn0*: 8/13**<br><br>Null allelles: 5/13 |
| *H. numata ethra* | Reserva Serra Bonita | BH16-0021 | ~500 | ~380 | |
| *H. numata ethra* | Reserva Serra Bonita | BH16-0022 | Null | Null | |
| *H. numata ethra* | Reserva Serra Bonita | BH16-0023 | ~500 | ~380 | |
| *H. numata ethra* | Reserva Serra Bonita | BH16-0024 | ~500 | ~380 | |
| *H. numata ethra* | Reserva Serra Bonita | BH16-0026 | Null | Null | |
| *H. numata ethra* | Reserva Serra Bonita | BH16-0027 | ~500 | ~380 | |
| *H. numata ethra* | Reserva Serra Bonita | BH16-0028 | Null | Null | |
| *H. numata ethra* | Reserva Serra Bonita | BH16-0029 | Null | Null | |
| *H. numata ethra* | Reserva Serra Bonita | BH16-0030 | ~500 | ~380 | |
| *H. numata ethra* | Reserva Serra Bonita | BH16-0031 | ~500 | ~380 | |
| *H. numata ethra* | Reserva Serra Bonita | BH16-0032 | ~500 | ~380 | |
| *H. numata ethra* | Reserva Serra Bonita | BH16-0033 | ~500 | ~380 | |
| *H. numata robigus* | Praia do Sono | BH16-0042 | ~500 | ~380 | **Arrangement *Hn0*: 11/11** |
| *H. numata robigus* | Praia do Sono | BH16-0043 | ~500 | ~380 | |
| *H. numata robigus* | Praia do Sono | BH16-0044 | ~500 | ~380 | |
| *H. numata robigus* | Praia do Sono | BH16-0045 | ~500 | ~380 | |
| *H. numata robigus* | Praia do Sono | BH16-0046 | ~500 | ~380 | |
| *H. numata robigus* | Praia do Sono | BH16-0047 | ~500 | ~380 | |
| *H. numata robigus* | Praia do Sono | BH16-0048 | ~500 | ~380 | |
| *H. numata robigus* | Praia do Sono | BH16-0049 | ~500 | ~380 | |
| *H. numata robigus* | Praia do Sono | BH16-0050 | ~500 | ~380 | |
| *H. numata robigus* | Praia do Sono | BH16-0051 | ~500 | ~380 | |
| *H. numata robigus* | Praia do Sono | BH16-0052 | ~500 | ~380 | |
| *H. numata silvana* | Oriximiná | HEL069 | ~500 | ~380 | **Arrangement *Hn0*: 2/2** |
| *H. numata silvana* | Oriximiná | HEL085 | ~500 | ~380 | |

**Table S3 | Sample name and metadata.** Includes sample id, species identification, morph, origin, sequencing depth, sample accession and type of analyses in which a given sample was used.

**See attached xls sheet file**

**Table S4 | Fst and Dxy between morphs or populations of *Heliconius* species and distance in km.** The first column represents the pair of populations compared. In some cases, where the subspecies ranges do not overlap (geographic races), the names of the subspecies studied are indicated instead of the populations. This is the case for *zuleika, melicerta, clearei* and “Peru” (geographic races/subspecies of *H. hecale*) for instance. Within *H. numata*, populations named Moyobamba, Tunel and Urahuasha are subsamples of the “Tarapoto” population indicated on Fig 1.

See next pages

| population1 | population2 | Species | $F_{ST}$ | Distance |
|---|---|---|---|---|
| Urahuasha | Moyobamba | H.numata (Amazon) | 0,018154 | 83 |
| Moyobamba | Michaela | H.numata (Amazon) | 0,021686 | 84 |
| Tunel | Moyobamba | H.numata (Amazon) | 0,01771 | 87 |
| Moyobamba | FG | H.numata (Amazon) | 0,055996 | 2924 |
| Moyobamba | Colombia | H.numata (Amazon) | 0,02706 | 879 |
| Moyobamba | Bresil | H.numata (Brazil) vs H.numata (Amazon) | 0,57124 | 3954 |
| Moyobamba | Ecuador | H.numata (Amazon) | 0,023722 | 550 |
| Moyobamba | Madre_de_Dios | H.numata (Amazon) | 0,049239 | 1042 |
| PuertoBermudez | Moyobamba | H.numata (Amazon) | 0,022862 | 524 |
| Moyobamba | La_Merced | H.numata (Amazon) | 0,028 | 574 |
| Urahuasha | Michaela | H.numata (Amazon) | 0,010698 | 55 |
| Urahuasha | Tunel | H.numata (Amazon) | 0,0040785 | 22 |
| Urahuasha | FG | H.numata (Amazon) | 0,038642 | 2873 |
| Urahuasha | Colombia | H.numata (Amazon) | 0,017033 | 911 |
| Urahuasha | Bresil | H.numata (Brazil) vs H.numata (Amazon) | 0,50322 | 3872 |
| Urahuasha | Ecuador | H.numata (Amazon) | 0,014587 | 606 |
| Urahuasha | Madre_de_Dios | H.numata (Amazon) | 0,035378 | 962 |
| Urahuasha | PuertoBermudez | H.numata (Amazon) | 0,0063779 | 458 |
| Urahuasha | La_Merced | H.numata (Amazon) | 0,017939 | 516 |
| Tunel | Michaela | H.numata (Amazon) | 0,0090059 | 53 |
| Michaela | FG | H.numata (Amazon) | 0,037372 | 2842 |
| Michaela | Colombia | H.numata (Amazon) | 0,012697 | 857 |
| Michaela | Bresil | H.numata (Brazil) vs H.numata (Amazon) | 0,53097 | 3891 |
| Michaela | Ecuador | H.numata (Amazon) | 0,010968 | 558 |
| Michaela | Madre_de_Dios | H.numata (Amazon) | 0,035196 | 993 |
| PuertoBermudez | Michaela | H.numata (Amazon) | 0,0069017 | 506 |
| Michaela | La_Merced | H.numata (Amazon) | 0,01899 | 567 |
| Tunel | FG | H.numata (Amazon) | 0,040786 | 2868 |
| Tunel | Colombia | H.numata (Amazon) | 0,01503 | 910 |
| Tunel | Bresil | H.numata (Brazil) vs H.numata (Amazon) | 0,56533 | 3867 |
| Tunel | Ecuador | H.numata (Amazon) | 0,013215 | 607 |
| Tunel | Madre_de_Dios | H.numata (Amazon) | 0,03683 | 958 |
| Tunel | PuertoBermudez | H.numata (Amazon) | 0,0047838 | 457 |
| Tunel | La_Merced | H.numata (Amazon) | 0,017184 | 515 |
| FG | Colombia | H.numata (Amazon) | 0,038624 | 2563 |
| FG | Bresil | H.numata (Brazil) vs H.numata (Amazon) | 0,59106 | 3143 |
| FG | Ecuador | H.numata (Amazon) | 0,040995 | 2857 |
| Madre_de_Dios | FG | H.numata (Amazon) | 0,064761 | 2661 |
| PuertoBermudez | FG | H.numata (Amazon) | 0,046453 | 2940 |
| La_Merced | FG | H.numata (Amazon) | 0,050235 | 3019 |
| Colombia | Bresil | H.numata (Brazil) vs H.numata (Amazon) | 0,53668 | 4353 |
| Ecuador | Colombia | H.numata (Amazon) | 0,0058787 | 407 |
| Madre_de_Dios | Colombia | H.numata (Amazon) | 0,039283 | 1697 |
| PuertoBermudez | Colombia | H.numata (Amazon) | 0,012295 | 1343 |
| La_Merced | Colombia | H.numata (Amazon) | 0,024572 | 1414 |
| Ecuador | Bresil | H.numata (Brazil) vs H.numata (Amazon) | 0,53699 | 4339 |
| Madre_de_Dios | Bresil | H.numata (Brazil) vs H.numata (Amazon) | 0,57678 | 2942 |
| PuertoBermudez | Bresil | H.numata (Brazil) vs H.numata (Amazon) | 0,64886 | 3525 |
| La_Merced | Bresil | H.numata (Brazil) vs H.numata (Amazon) | 0,57264 | 3535 |
| Madre_de_Dios | Ecuador | H.numata (Amazon) | 0,038934 | 1521 |
| PuertoBermudez | Ecuador | H.numata (Amazon) | 0,012336 | 1064 |
| La_Merced | Ecuador | H.numata (Amazon) | 0,021665 | 1121 |

**Table S4 (suite)**

| population1 | population2 | Species | $F_{ST}$ | Distance |
|---|---|---|---|---|
| PuertoBermudez | Madre_de_Dios | H.numata (Amazon) | 0,03854 | 585 |
| Madre_de_Dios | La_Merced | H.numata (Amazon) | 0,043319 | 603 |
| PuertoBermudez | La_Merced | H.numata (Amazon) | 0,015199 | 87 |
| amaryllis | aglaope | H.melpomene | -4,84E-05 | 30 |
| Ser | But | H.pardalinus | 0,42796 | 30 |
| zuleika | melicerta | H.hecale | 0,031355 | 400 |
| Peru | But | H.pardalinus | 0,010075 | 400 |
| Ser | Peru | H.pardalinus | 0,42679 | 430 |
| Clarescens | boulleti | H.ismenius | -0,015628 | 511 |
| Peru | Ecuador | H.elevatus | 0,012688 | 571 |
| Peru | melicerta | H.hecale | 0,29052 | 1776 |
| melicerta | clearei | H.hecale | 0,35583 | 1900 |
| zuleika | Peru | H.hecale | 0,31119 | 2100 |
| Peru | clearei | H.hecale | 0,39312 | 2230 |
| zuleika | clearei | H.hecale | 0,37628 | 2300 |
| French_Guiana | Ecuador | H.elevatus | 0,20174 | 2730 |
| Peru | French_Guiana | H.elevatus | 0,19544 | 2900 |
| French_Guiana | amaryllis | H.melpomene | 0,33315 | 2900 |
| French_Guiana | aglaope | H.melpomene | 0,30753 | 2930 |
| Peru | Brazil | H.ethilla | 0,43403 | 4150 |
| Tunel | SE_Brazil | H.numata (SE Brazil) vs H.numata (Amazon) | 0,48327 | 4107 |
| Urahuasha | SE_Brazil | H.numata (SE Brazil) vs H.numata (Amazon) | 0,43968 | 4112 |
| SE_Brazil | Bresil | H.numata (Brazil) vs H.numata (SE Brazil) | 0,012293 | 862 |
| SE_Brazil | Colombia | H.numata (SE Brazil) vs H.numata (Amazon) | 0,46257 | 4101 |
| SE_Brazil | Ecuador | H.numata (SE Brazil) vs H.numata (Amazon) | 0,46496 | 4307 |
| SE_Brazil | FG | H.numata (SE Brazil) vs H.numata (Amazon) | 0,51405 | 2074 |
| SE_Brazil | La_Merced | H.numata (SE Brazil) vs H.numata (Amazon) | 0,49424 | 3991 |
| SE_Brazil | Madre_de_Dios | H.numata (SE Brazil) vs H.numata (Amazon) | 0,49896 | 3397 |
| SE_Brazil | Michaela | H.numata (SE Brazil) vs H.numata (Amazon) | 0,45999 | 4103 |
| SE_Brazil | Moyobamba | H.numata (SE Brazil) vs H.numata (Amazon) | 0,49207 | 4185 |
| SE_Brazil | PuertoBermudez | H.numata (SE Brazil) vs H.numata (Amazon) | 0,5753 | 3946 |

**Table S5 | Effective population sizes and divergence times in years as inferred by G-PhoCS.**
Historical effective population size (Ne) and split time (T) are referred to using species initials; for instance, the population size of the ancestror of *H. pardalinus* and *H. elevatus* is referred to as Ne_PE. Mean, standard deviation and HPD interval (minimum and maximum) at 95% are presented. NumataA stands for *H. numata* from the Amazon and French Guiana, NumataB stands for *H. numata* from the Atlantic forests of Brazil.

| Parameter | Mean | S.D. | HPDmin | HPDmax |
|---|---|---|---|---|
| **Ne_Cydno** | 5311544 | 57966.764 | 5195578.9 | 5423236.8 |
| **Ne_NumataA** | 23089618 | 1089100.2 | 20680171 | 25441513 |
| **Ne_NumataB** | 1144291 | 15608.681 | 1114302.6 | 1175578.9 |
| **Ne_Pardalinus** | 15132304 | 972850.99 | 12661145 | 16301789 |
| **Ne_Elevatus** | 11833709 | 805558.7 | 10600079 | 13721697 |
| **Ne_Ismenius** | 1023584.4 | 11708.262 | 1000789.5 | 1046605.3 |
| **Ne_PE** | 6835145.4 | 498638.94 | 6224605.3 | 8051000 |
| **Ne_Numata** | 4963871.5 | 431830.73 | 4406328.9 | 5621407.9 |
| **Ne_IN** | 3492728.6 | 1567408.1 | 628144.74 | 6529223.7 |
| **Ne_PEIN** | 2441101.7 | 1818552.5 | 74986.842 | 5999723.7 |
| **Ne_CPEIN** | 3199022.4 | 39979.576 | 3121052.6 | 3278289.5 |
| **T_PE** | 1375611.6 | 84100.673 | 1171210.5 | 1452960.5 |
| **T_Numata** | 1429080.5 | 48163.774 | 1348973.7 | 1541921.1 |
| **T_IN** | 2412324.2 | 187762.79 | 1903421.1 | 2569671.1 |
| **T_PEIN** | 2543031.1 | 8980.2061 | 2525486.8 | 2560947.4 |
| **T_CPEIN** | 2543227.2 | 8977.8798 | 2525500 | 2560973.7 |

**Table S6 | Migration bands analysed with G-PhoCS.**
Mean values, standard deviation, minimum and maximum confidence interval at 95% showing the probability that the estimated total migration was greater than 0.

| | Mean | S.D. | HPD.C.I..minimum | HPD.C.I..maximum | Probability |
|---|---|---|---|---|---|
| *Numata -> Ismenius* | 0.80433 | 0.22333 | 0.49155 | 1.10236 | 1 |
| *Ismenius -> Numata* | 0.00017 | 0.00048 | 0 | 0.00095 | 0 |
| *Elevatus -> Pardalinus* | 1.77953 | 0.53181 | 0.65674 | 2.41723 | 1 |
| *Pardalinus -> Elevatus* | 0.0567 | 0.10494 | 0 | 0.32894 | 0.282 |
| *PE -> IN* | 0.00019 | 0.00057 | 0 | 0.00113 | 0 |
| *IN -> PE* | 0.00015 | 0.00047 | 0 | 0.00092 | 0 |
| *PEIN -> Cydno* | 0.00039 | 0.00161 | 0 | 0.00224 | 0 |
| *Cydno -> PEIN* | 8e-05 | 0.00036 | 0 | 0.00034 | 0 |
| *Numata -> PE* | 0.19306 | 0.04095 | 0.12593 | 0.25093 | 1 |
| *PE -> Numata* | 0.02067 | 0.02045 | 0 | 0.05907 | 0.342 |
| *Ismenius -> PE* | 0.00018 | 0.00044 | 0 | 0.00099 | 0 |
| *PE -> Ismenius* | 0.11323 | 0.02506 | 0.06787 | 0.16708 | 1 |
| *Pardalinus -> Ismenius* | 0.00033 | 0.00084 | 0 | 0.00189 | 0 |
| *Ismenius -> Pardalinus* | 0.00013 | 0.00025 | 0 | 0.00066 | 0 |
| *Elevatus -> Ismenius* | 0.00087 | 0.00202 | 0 | 0.00609 | 0 |
| *Ismenius -> Elevatus* | 9e-05 | 0.00016 | 0 | 0.00045 | 0 |

**Table S7: Model choice from ∂a∂i.** Comparison between *H. numata* from peru and *H. numata* from Brazil and *H. numata* from Peru versus *H. pardalinus.* SI = Strict Isolation, IM = Isolation W. Migration, AM = Ancient Migration. G = suffix indicating Growth in the daughter populations. 2N = suffix indicating heterogeneity of effective population size. A = Suffix indicating growth in the Ancestral population.

| ***H. numata* Peru (Amazonian) – vs – *H. numata* Brasil (Mata Atlântica)** | | ***H. Numata* Peru (Amazonian) – vs – *H. pardalinus*** | |
|---|---|---|---|
| **Model** | **AIC** | **Model** | **AIC** |
| SI2NG | 13878 | SI2N | 10297 |
| SI2N | 11405 | SI2NG | 9917 |
| AM2N | 9838 | IM2N | 6211 |
| IM2N | 9762 | AM2N | 6183 |
| SC2N | 8963 | SC2N | 5923 |
| IM2NG | 7587 | SC2NG | 4029 |
| AMA2NG | 6713 | SCA2NG | 4029 |
| IMA2NG | 6179 | SIA2NG | 3310 |
| AM2NG | 6089 | IM2NG | 3131 |
| SC2NG | 5542 | AMA2NG | 2932 |
| ***SCA2NG*** | ***2888*** | ***IMA2NG*** | ***2099*** |

**Text S1 | Supplementary Methods**

*Analysis of the supergene occurrence Atlantic populations of Brazil of H. numata*

*Heliconius numata robigus* and *H. n. ethra* are distributed along the South-Eastern coastal border of Brazil (Figure S2). They are both confined to the Atlantic forests of Brazil (*Mata Atlântica* biome) and are geographically isolated from the rest of the races of *H. numata,* which are predominantly distributed throughout the Amazon basin (grey area in Figure S2). Populations of *H. n. robigus* and *H. n. ethra* are not locally polymorphic (Brown and Mielke, 1972), something we confirmed by observing the collected specimens in two of the biggest Lepidoptera collections in Brazil, that of the Museo de Zoologia da Universidade de São Paulo (MZ/USP) and that of the Museo Nacional da Universidade Federal de Rio de Janeiro (MN/UFRJ). Instead, *H. n. robigus* and *H. n. ethra* share a quite large transition zone where any of these races or wing color intermediates between them can be found. Importantly, however, these races are distinguishable on the basis of rather few wing colour elements (Figure S2) when compared to the large phenotypic differences existing between polymorphic morphs of *H. numata*. We collected 11 and 13 specimens of *H. numata* forms *robigus* and *ethra* at Praia do Sono, Paraty, Rio de Janeiro State (see red circle in Figure S2) and at Reserva Serra Bonita, Camacã, Bahia State (see orange circle in Figure S2), respectively. Butterfly bodies were preserved in a NaCl-saturated EDTA-DMSO solution and wings were stored separately in glassine envelopes. We isolated gDNA from the preserved bodies using the DNeasy Blood and Tissue Kit (Qiagen). We genotyped gene *cortex* for the collected *H. n. robigus* and *H. n. ethra* specimens to determine whether or not the Brazilian populations harbour the ancestral *Hn0* gene order at the supergene *P. Cortex* controls whole-wing variation in black, yellow, white, and orange/red elements in *H. numata* color pattern (Nadeau *et al.* 2016). Allelic variants at gene *cortex* have been consistently associated with the different gene orders at the supergene *P* in Peruvian populations of this species (Joron et al., 2011). Primers that allow assessing this allelic variation are anchored on exons 2 and 3 of gene *cortex* (Table S1). Standard PCR master mixes and thermal cycling conditions were used. Thermal cycling conditions were 94°C for 3 min, 40 cycles of 94°C for 30 sec, 53°C annealing for 45 sec, 72°C for 90 sec, and a final extension at 72°C for 5 min. We scored amplicon length variation of gene *cortex* in agarose gels stained with BEt. Two specimens of *H. numata silvana* from an Amazonian population of Brazil were used as a control for the size of the *silvana*-like band (arrangement *Hn0*). Results (Table S2) show that *H. numata* populations from the Atlantic forest of Brazil are devoid of any rearrangements.

*Analyses of the slope of Fst versus distance, as measured in kilometers*

As stated in the main text, we tested for the existance and intensity of an isolation by distance (IBD) signal among species and between population of *H. numata* using a linear model. If IBD is stronger in species not polymorphic for the inversion we should observed significantly steeper slopes in these species. The slopes of $F_{ST}$/1-Fst versus log(distance) in km was calculated using the R package *lsmeans* (Lenth 2016); the slope difference among species or between populations within species was estimated with an ANOVA and its significance evaluated with function pairs of this package.

As shown in Fig. 2, *H. numata* has a much lower slope except for those Fst which involved the Brazilian samples. In order to quantify whether these slopes were significantly different, we performed the following analyses, using ANOVA and the package *lsmeans*. We analysed the slope of Fst/1-Fst vs. log10(distance) involving only *H. numata* and compared with the slope of Fst vs. distance within all other *Heliconius* (i.e., *H. elevatus*, *H. ethilla*, *H. ismenius*, *H. hecale*, *H. pardalinus* and *H. melpomene*) together, and tested whether these slopes were significantly different. Using the function lm of the package *lsmeans*, we analysed the interaction of distance and group (all other Heliconii or *H. numata* only) on Fst. The analysis of variance (ANOVA) showed that both distance and group have a significant effect on Fst, but not their interaction. Using the functions *lm* and pairs of *lsmeans*, we observed that the slope of Fst vs. distance for each group were significantly different (p=0.04) but with a greater slope for *H. numata*. We proceeded then to a similar analysis removing the Brazilian comparisons from *H. numata* set. In that case, the analysis of variance of Fst showed that distance, species and the interaction between distance and group had a significant effect. Using *lstrends*, the difference between the slope of Fst vs. distance for *H. numata* and other *Heliconius* was significant (p<0.0042).

*Testing for an effect of the inversion on population differentiation*

In order to test whether the difference in slopes in *H. numata* came from the absence of the inversion supergene in the Brazilian samples, we performed similar analyses to those above mentioned, with and without chromosome 15, where the inversions are located. Firstly, we tested the significance of the slope of Fst/(1-Fst) vs. log(distance) using an analysis of variance of a linear model including as response variable distance, group (Amazonian or Atlantic) and their interaction. All these three components were significant. Using *lstrends* and the *pairs* function, the slope of Fst vs. distance for *H. numata* without the Atlantic locality was significantly different from the slope of Fst vs. distance for comparisons which included the Atlantic population ($p<0.0001$). Therefore the slope of Fst vs distance was significantly higher when including the Atlantic population.

We then performed the same analysis for Fst computed without chromosome 15. The analysis of variance resulted in significant effects on Fst from distance, group (Atlantic or Amazon) and the interaction of distance and group. The slope of Fst vs. distance for comparisons including only Amazonian populations and the slope including all comparisons were also significantly different ($p<0.0001$). Therefore the slope of Fst vs distance without chromosome 15 was significantly higher when including the Atlantic population. The result were the same with or without chromosome 15 and suggests that including or not the inversion in our computation did not have a major influence. Overall the slope of the signal of IBD was lower when comparing only Amazonian populations than when the Atlantic population was considered.
